# Common proanthocyanidin-rich foods modulate gastrointestinal blooms of *Akkermansia muciniphila* in a diet-dependent manner

**DOI:** 10.1101/2021.11.07.466338

**Authors:** Katia S. Chadaideh, Kevin E. Eappen, Brandi E. Moore, Rachel N. Carmody

**Affiliations:** Department of Human Evolutionary Biology, Harvard University, 11 Divinity Avenue, Cambridge, MA 02138, USA

## Abstract

Developing methods to modulate growth of the mucin-degrading gut bacterium *Akkermansia muciniphila* could benefit patients with different health needs, as *A. muciniphila* has been associated with both positive metabolic health outcomes and detrimental neurodegenerative outcomes. Growth of *A. muciniphila* is sensitive to plant-derived polyphenols, and particularly proanthocyanidins (PACs), when administered in isolated form at supraphysiological doses. However, it remains unclear whether doses sufficient for these effects are achievable via diet. Here, we explore the extent to which nutritionally relevant doses of common polyphenol-rich foods – berries, wine, and coffee – influence *A. muciniphila* abundance in C57BL/6J mice under varying dietary conditions. By administering polyphenol-rich whole foods, comparing polyphenol-depleted and PAC-rich versus PAC-poor food supplements, and through gradient PAC-dosing experiments, we show that PAC-rich foods uniquely induce *A. muciniphila* growth at doses that are feasibly achieved through routine diet. Notably, the effects of PAC supplementation were detected against a high-fat diet but not a low-fat control diet background, highlighting the importance of habitual diet strategies in either amplifying or mitigating the prebiotic effects of PAC-rich food consumption. Ultimately, our work suggests that both PACs and diet influence *A. muciniphila* abundance with downstream impacts for human health.

## Introduction

Given compelling links between the gut microbiota and numerous aspects of human health, there is great potential in developing therapies that target the gut microbial community. Although the gut microbiome and its interactions with host physiology are highly sensitive to diet (Carmody et al., 2015; Chadaideh and Carmody, 2021; David et al., 2014; Sonnenburg et al., 2016; Turnbaugh et al., 2010), the complex network of microbial relationships in the gut often results in interventions having unpredictable or difficult-to-replicate outcomes. Therefore, there is a pressing need to identify interventions that lead to predicable increases or decreases in specific microbial taxa implicated as causal agents in specific health phenotypes.

One candidate intervention targeting the gut microbiota is the administration of diet-derived polyphenols, which have been associated with lowered risks of chronic metabolic and inflammatory diseases (Del Bo et al., 2019; Domínguez-Avila et al., 2020; Halliwell, 2007). Specifically, the subclass of proanthocyanidins (PACs), which are responsible for the bitter taste of berries and found in wine and other tannin-rich foods (Gu et al., 2004), have been strongly associated with attenuating obesity, insulin resistance, and systemic inflammation in mice (Anhê et al., 2015; Dixon et al., 2004; Roopchand et al., 2015; Zhang et al., 2018), as well as reducing cardiovascular disease and diabetes risk in humans (Blumberg et al., 2013; Del Bo et al., 2019; Gu et al., 2004).

The mechanisms by which polyphenols confer beneficial metabolic effects are unclear, but their role in cardiometabolic protection may at least partially derive from changes in the gut microbiota, and specifically, increased abundance of the bacterial species *Akkermansia muciniphila* (Anhê et al., 2015; Everard et al., 2013; Kim et al., 2020; Régnier et al., 2020; Roopchand et al., 2015). In studies investigating polyphenol supplementation in mice, administration of isolated PACs at levels representing 1% of the diet by weight was shown to increase the relative abundance of *A. muciniphila* from <2% to >40% within days (Anhê et al., 2015; Roopchand et al., 2015). Interestingly, *A. muciniphila* blooms in mice preceded changes in host gene expression relevant to low-grade systemic inflammation (Everard et al., 2013; Zhang et al., 2018), and correlated with reduced adiposity and insulin insensitivity on a high-fat diet (Anhê et al., 2015; Roopchand et al., 2015; Zhang et al., 2018). *A. muciniphila* enrichment was also recently shown to improve metabolic health in a randomized double-blind placebo-controlled study in overweight/obese insulin-resistant human subjects (Depommier et al., 2019). And while the specific mechanisms though which polyphenol-induced enrichment of *A. muciniphila* improves metabolic health also remain unknown, *A. muciniphila* secretes the protein P9, which was recently found to be sufficient to induce glucagon-like peptide-1 (GLP-1) secretion and brown fat thermogenesis in mice fed a high-fat diet (Yoon et al., 2021).

Despite evidence that *A. muciniphila* enrichment improves metabolic health, increased abundance of this microbe does not appear to be universally advantageous. For example, investigations into the role of the gut-brain axis in neurological function have revealed correlations between *A. muciniphila* and neurodegenerative disease (Fang et al., 2020). Recent studies investigating the gut microbiota of Parkinson’s patients demonstrated a link between increased relative abundance of the genus *Akkermansia* and a progression of symptoms (Haikal et al., 2019; Heintz-Buschart et al., 2018; Romano et al., 2021; Scheperjans et al., 2015; Vidal-Martinez et al., 2020), and *A. muciniphila* has been suggested as a biomarker for diagnosing Parkinson’s patients relative to healthy age-matched controls (Bedarf et al., 2017). One explanation for the diverging roles of *A. muciniphila* may be that the genomic diversity of *A. muciniphila* has yet to be fully characterized. Recent work using metagenome-assembled genomes (MAGs) has identified distinct subspecies-level genomic variation in *A. muciniphila* that may be associated with differences in functional capacity (Karcher et al., 2021). Furthermore, *A. muciniphila* has been shown to induce altered host immune responses depending on broader gut microbial community composition in mice (Ansaldo et al., 2019), and this context-dependence has been proposed as a reason for varying host health outcomes associated with *A. muciniphila* enrichment (Haikal et al., 2019).

Given links between *A. muciniphila* and divergent metabolic and neurodegenerative outcomes, there is great therapeutic potential in developing methods to modulate *A. muciniphila* abundance within the gut. Previous studies in mice have reported strong *A. muciniphila* blooms in response to polyphenol administration (Anhê et al., 2017; Régnier et al., 2020; Roopchand et al., 2015; Zhang et al., 2018), but their experimental designs used purified compounds divorced from their whole food form. Isolated phenolic compounds are volatile and degrade easily (Volf et al., 2014), making them difficult and costly to administer commercially. Furthermore, although polyphenols are found in a wide range of foods commonly consumed in industrialized populations (Manach et al., 2004), the doses previously tested in mice have been 20-50 times greater, after allometric scaling, than what would typically be consumed by humans in industrialized settings (Del Bo et al., 2019). Therefore, it remains unclear whether ingesting polyphenols at levels achievable via foods already consumed in an industrialized diet could similarly foster blooms of *A. muciniphila*. In addition, to our knowledge, studies exploring how to selectively impede *A. muciniphila* growth have yet to be performed. Understanding how environmental drivers like diet can shape gut microbiota dynamics that influence *A. muciniphila* growth could advance new avenues to mitigate metabolic and neurodegenerative disease via the gut microbiota.

In this study, we explore the extent to which consuming nutritionally relevant doses of common polyphenol-rich foods influences the growth of *A. muciniphila* in mice under high-fat versus low-fat dietary conditions. By feeding native and polyphenol-depleted versions of foods that vary in their underlying composition of polyphenolic compounds, we show that the modulation of *A. muciniphila* is driven not by total polyphenolic content, but rather by PAC content. By observing changes in the gut microbiome under PAC-rich versus PAC-poor conditions, and through gradient PAC-dosing experiments, we show that PAC-rich foods commonly consumed in industrialized settings uniquely induce *A. muciniphila* growth at doses that can be achieved through routine diet. Notably, the effects of PAC supplementation were detected against a high-fat but not a low-fat diet background, highlighting that the prebiotic effects of PACs may depend on the broader diet strategy. Overall, our findings indicate that interactions between PAC-rich food supplementation and overall diet composition may play a significant role in modulating *A. muciniphila* growth.

## Results

### Dried blueberry consumption leads to rapid blooms of *A. muciniphila*

To date, studies supporting an association between polyphenol consumption and blooms of *A. muciniphila* have relied exclusively on purified polyphenol extracts (Anhê et al., 2015, 2016, 2017; Régnier et al., 2020; Roopchand et al., 2015; Zhang et al., 2018). Therefore, our first goal was to establish whether polyphenols administered in their whole food matrix could likewise induce *A. muciniphila* growth. To do this, we first examined changes in gut microbial community composition among mice reared for 5 days on homogeneous *ad libitum* diets of polyphenol-rich dried grapes (raisins), dried cranberries, or dried blueberries relative to a standard chow control group (**Figure 1A**). Each of these berries is rich in polyphenolic compounds although they differ in total polyphenolic abundance and composition (**Figure 1B-C**), and each berry has been associated with metabolic benefits in humans (Curtis et al., 2019; Fulgoni et al., 2017; Pourmasoumi et al., 2020; Stote et al., 2020; Wijayabahu et al., 2019).

**Figure 1:**
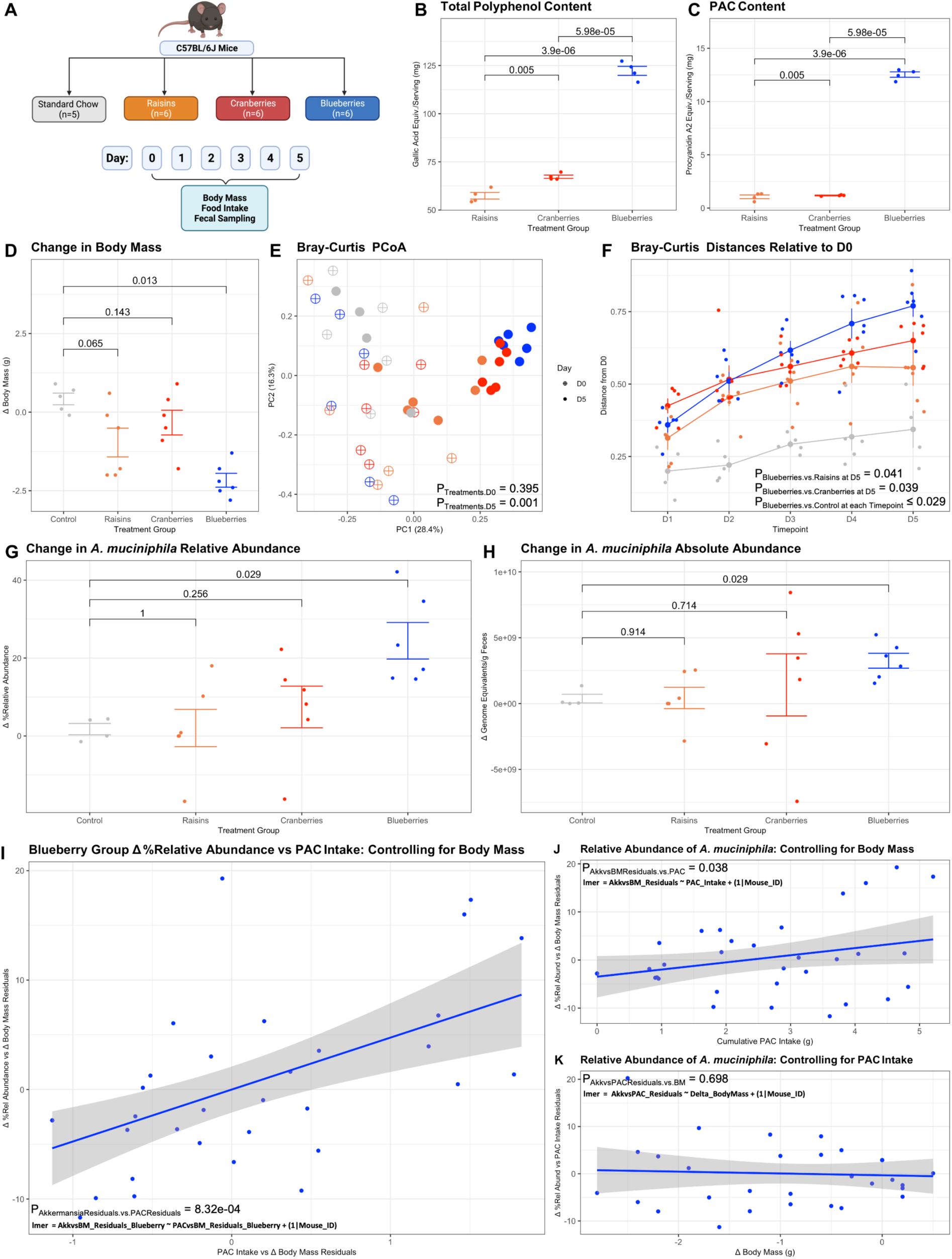
Dried blueberry consumption results in significant differences in body mass and gut microbial community structure in mice. **(A)** We investigated the effects of dried berry consumption (blueberries, cranberries, and raisins) on host and microbial phenotypic effects in adult male C57BL/6J mice (n=6) compared to consumption of a standard chow control (n=5) over a 5-day dietary intervention. **(B)** Total polyphenol content (TPC) contained in a serving of each dried berry source (1/4 cup), measured using the Folin-Ciocalteau assay. **(C)** Proanthocyanidin (PAC) content per serving of dried berry source (1/4 cup), measured using the DMAC assay. **(D)** Change in body mass from Day 0 to Day 5. **(E)** Bray-Curtis PCoA plot for samples collected on Day 0 versus Day 5. Association of samples by treatment group determined at each timepoint using PERMANOVA. **(F)** Bray-Curtis distances of fecal samples at each timepoint for each mouse relative to its own baseline sample. **(G)** Change in relative abundance of *A. muciniphila* from Day 0 to Day 5. **(H)** Change in absolute abundance of *A. muciniphila* from Day 0 to Day 5. **(I)** Comparing the change in relative abundance of *A. muciniphila* in response to cumulative PAC intake and when controlling for their relationships to body mass. **(J)** The effect of cumulative PAC intake on *A. muciniphila* relative abundance when controlling for body mass. **(K)** The effect of changes in body mass on *A. muciniphila* relative abundance when controlling for PAC intake. **(B-C)** Statistics determined using student’s t-test and Benjamini-Hochberg correction for false discovery rate. **(D, F-H)** Statistics determined using Mann-Whitney U test with Benjamini-Hochberg correction for false discovery rate. **(I-K)** Statistics determined using linear mixed effects model.

The differing total polyphenol content (TPC) and proanthocyanidin (PAC) content of the raisins, cranberries, and blueberries used in this study (**Figure 1B-C**) allowed us to evaluate any differential phenotypic effects of polyphenol intake. Consumption of any of the berry treatments led to significantly lower mRNA expression in the proximal colon of markers of gut inflammation (IL-6, IL-1β and TNFα), gut epithelial integrity (Muc2 and Ocln, but not Intectin), and, to some degree, antimicrobial peptides (Reg3γ and Pla2g2a) (**Figure S1F-M**). However, by the end of the 5-day intervention, mice consuming blueberries (highest TPC and PAC content) exhibited a significant decrease in total body mass compared to controls (**Figure 1D**), an effect not observed in mice consuming raisins (lowest TPC and PAC content) or cranberries (intermediate TPC and low PAC content). Notably, accelerated loss of body mass was not driven by reduced food consumption in the blueberry group, as food consumption was comparable across groups (p=0.356, Kruskal-Wallis). Rather, change in body mass per unit food intake decreased significantly among mice consuming blueberries (p=0.002, linear mixed effects model, **Figure S1A**), while the relationship between change in body mass and total food intake among mice consuming either raisins or cranberries did not significantly differ from controls (**Figure S1A**). Foreshadowing the possibility that PACs exert unique effects, we also detected significant positive longitudinal relationships between decreases in total body mass and total PAC intake (p=4.92e-16, linear mixed effects model) among mice in all three berry treatment groups.

Gut microbial community signatures of mice were also distinct by treatment group at endpoint (p=0.001, pseudo-F=4.97, PERMANOVA; **Figure 1E**). Blueberry consumption uniquely resulted in significant changes in Bray-Curtis distances relative to baseline when compared to the control diet at every day of the study, as well as when compared to either cranberries or raisins by Day 5 (**Figure 1F**). Changes in whole community composition were not an artifact of changes in alpha-diversity (Shannon index), the ratio of Firmicutes to Bacteroidetes relative abundance as an index of tradeoffs between the two dominant bacterial phyla in the gut, or total bacterial abundance as assessed by universal 16S qPCR (**Figure S1C-E**). However, increased PAC intake did significantly correspond with longitudinal increases in the relative abundance of *A. muciniphila* (p=6.94e-14, linear mixed effects model). Correspondingly, blueberry consumption correlated with striking blooms in both the relative and absolute abundance of *A. muciniphila* compared to the control group (**Figure 1G-H**), including after controlling for cumulative food intake (p=0.0104, linear mixed effects model; **Figure S1B**).

Because mice on the blueberry treatment exhibited strong correlations between *A. muciniphila* relative abundance, PAC intake, and changes in body mass, we examined the degree to which *A. muciniphila* enrichment was the result of PAC intake versus an artifact of weight loss. When controlling for their associations with body mass, *A. muciniphila* relative abundance maintained a significant positive relationship with cumulative PAC intake (p=8.32e-04, linear mixed effects model, **Figure 1I**). Furthermore, whereas PAC intake maintained a positive effect on *A. muciniphila* relative abundance when controlling for its relationship to body mass (p=0.038, linear mixed effects model, **Figure 1J**), body mass exhibited no effect on *A. muciniphila* when controlling for its relationship to PAC intake (p=0.698, linear mixed effects model, **Figure 1K**). These results suggest that PAC intake, not weight loss, is the primary driver of *A. muciniphila* enrichment among mice consuming dried blueberries.

### *A. muciniphila* growth promoted by PAC-rich foods, not generally polyphenol-rich foods, against the backdrop of a high-fat diet

Because blueberries were highest in TPC as well as PACs relative to the other berry groups (**Figure 1B-C**), we next proceeded to confirm that such effects were unique to PAC-rich as opposed to generally polyphenol-rich foods. To isolate gut microbial effects attributable to PACs versus total polyphenols and to investigate whether the broader diet shapes the response of *A. muciniphila* to foods rich in these phenolics, we gavaged mice with polyphenol-rich red wine or coffee, or polyphenol-depleted versions of the same red wine or coffee, as daily prebiotic supplements on top of either a high-fat diet (HFD) or a standard chow control diet (**Figure 2A**). We selected red wine and coffee for testing because they are similarly rich in TPC but differ significantly in their polyphenol composition (Gu et al., 2004), with wine being rich in PACs while coffee is not (**Figure 2B-C**). Supplementation was achieved by gavaging mice daily with 100µL/25g of a solution containing 1.47g dry weight/mL dealcoholized red wine, 1.30g dry weight/mL polyphenol-free dealcoholized wine (winePF), 0.606g dry weight/mL decaffeinated coffee, or 0.497g dry weight/mL polyphenol-free decaffeinated coffee (coffeePF), allometrically scaled doses equivalent to 750mL (i.e., 1 bottle of wine or 25 oz of coffee) for a 65kg human (**Figure 2A**). We compared each of these conditions to a vehicle control group, which was gavaged daily with 100µL/25g of nuclease-free water. Under the hypothesis that PAC consumption drives *A. muciniphila* abundance, we would expect increased *A. muciniphila* only among the wine treatment, as the other treatments are either experimentally depleted in PAC (winePF) or high intrinsically low in PACs (coffee or coffeePF).

**Figure 2:**
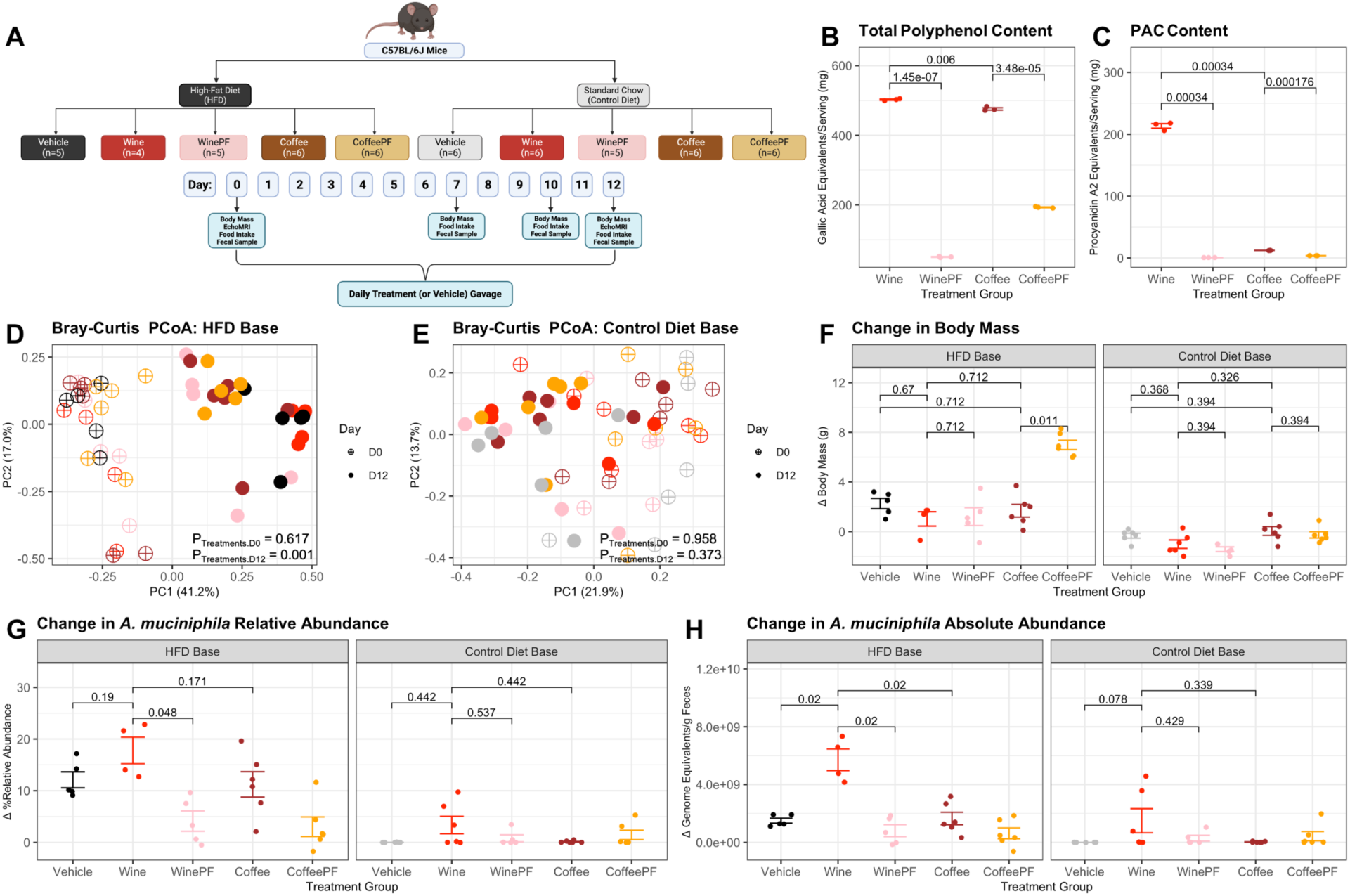
Wine versus coffee supplementation impacts microbial community structure under high-fat diet (HFD) but not control diet feeding in mice. **(A)** Adult male C57BL/6J mice consuming either a HFD or control diet base were administered one of the following prebiotic treatments over a 12-day dietary intervention: alcohol-free wine (wine), alcohol-free and polyphenol-free wine (winePF), caffeine-free coffee (coffee), caffeine-free and polyphenol-free coffee (coffeePF), or a vehicle control (n=4-6 per condition). Treatments or vehicle controls were administered daily via oral gavage. **(B)** Total polyphenol content (TPC) per recommended serving of each treatment (5 oz of wine product or 6oz of coffee product) as measured by the Folin-Ciocalteau assay. **(C)** Proanthocyanidin (PAC) content per recommended serving of each treatment (5 oz of wine product or 6oz of coffee product) as measured by the DMAC assay. **(D)** Bray-Curtis PCoA plot for HFD treatments across Day 0 versus Day 12 of the study. **(E)** Bray-Curtis PCoA plot for control diet treatments across Day 0 versus Day 12 of the study. **(F)** Change in body mass from Day 0 to Day 12 across treatment groups administered each diet base. **(G)** Change in relative abundance of the *A. muciniphila* from Day 0 to Day 12 across treatment groups administered each diet base. **(H)** Change in absolute abundance of *A. muciniphila* from Day 0 to Day 12 across treatment groups administered each diet base. **(B-C)** Statistics determined using Student’s t-test and Benjamini-Hochberg correction for false discovery rate. **(D-E)** Association of samples by treatment group determined at each timepoint using PERMANOVA. **(F-H)** Statistics determined using Mann-Whitney U test with Benjamini-Hochberg correction for false discovery rate.

As seen in prior work (Carmody et al., 2015; Turnbaugh et al., 2006), transitioning from standard chow to an HFD generally altered Bray-Curtis distances relative to baseline (**Figure S2B**), while mice consuming the control diet exhibited minimal changes relative to baseline (**Figure S2O**). Importantly, however, we observed distinct interaction effects between the diet base and prebiotic supplementation that impacted gut microbial phenotypes. Among HFD-fed mice, gut microbial community composition differed by prebiotic treatment by Day 12 of the study (p=0.001, pseudo-F=2.87, PERMANOVA; **Figure 2D**), but such differentiation was not observed among mice fed the control diet (p=0.373, pseudo-F=1.061, PERMANOVA; **Figure 2E**). The compositional changes with treatment among HFD-fed mice could not be attributed to differences in alpha-diversity (Shannon index), Firmicutes to Bacteroidetes relative abundance, or total bacterial abundance (**Figure S2C-E,P-R**), Instead, treatment differences seen on the HFD were driven in large part by changes in *A. muciniphila* abundance (p=1.38e-08, coef=5.568, MaAsLin2), with the HFD-wine supplementation inducing the greatest enrichment of *A. muciniphila* by endpoint (p=0.001, coef=9.063, MaAsLin2).

Consistent with our hypothesis, the HFD-wine group experienced the only significant changes in *A. muciniphila* abundance, with the HFD-wine group exhibiting increased *A. muciniphila* relative abundance compared to the HFD-winePF group (**Figure 2H**), as well as increased absolute abundance compared to the HFD-winePF, HFD-coffee, or HFD-vehicle groups (**Figure 2G**). As seen in the blueberry case (**Figure 1I-K**), enrichment of *A. muciniphila* in the HFD-wine group was not attributable to underlying changes in body mass, as change in body mass was similar among the HFD-wine, HFD-winePF, HFD-coffee, and HFD-vehicle groups (**Figure 2F**). That wine supplementation did not induce similar effects on *A. muciniphila* when administered under control diet feeding (**Figure 2G-H**), suggests that PAC-induced blooms of *A. muciniphila* may be enhanced by a diet high in fat and low in fiber.

Overall, we observed no significant effects of either wine or coffee supplementation on host body mass or adiposity relative to the vehicle control (**Figure 2F, Figure S2A,N**). Moreover, in contrast to signatures of proximal colon expression seen in the dried berry study (**Figure S1F-M**), wine and coffee treatments did not broadly alter expression of markers of inflammation or gut barrier integrity in the proximal colon (**Figure S2F-M,S-Z**), although we did observe a consistent increase in *Ocln* expression among both HFD-fed and control diet-fed mice on the wine treatment relative to winePF and coffee supplementation (**Figure S2J,W**). The sources of these experimental differences are unknown, but may be attributable to differences in effective dose between homogeneous (berry) and supplemented (coffee and wine) diets, physicochemical differences between solid and liquid diets, or differences in macronutrient composition across the test substrates. To discriminate among these variables, we next proceeded to test the dose of PACs required to stimulate *A. muciniphila* blooms under conditions that controlled for background diet.

### PACs induce *A. muciniphila* blooms at a distinct, diet-relevant dose in mice consuming high-fat diets, while mice consuming control diets exhibit a blunted response to PACs

Our polyphenol composition experiment confirmed that supplementing mice with PAC-rich red wine, but not PAC-depleted red wine or PAC-poor coffee, can increase *A. muciniphila* abundance regardless of changes in body composition, and that the effects of PAC intake on the murine gut microbiome may be enhanced against a HFD base. Although the dose of red wine we administered was selected to lie at the high end of human nutritional relevance (the allometric equivalent of 1 bottle of wine for humans, or 1.47g dry weight/mL per 25g mouse), we next sought to determine whether blooms in *A. muciniphila* could occur at lower levels of PAC intake under a HFD and to validate our prior finding that the prebiotic effect of PACs is relatively minimal on a control diet. To accomplish this, we performed a PAC dosing experiment using the same purified grape-derived PACs used in previous studies (Roopchand et al., 2015; Zhang et al., 2018). We selected the following range of PAC doses to administer to mice consuming either HFD or control diet bases: (1) a high PAC concentration (9mg/25g, or 9PAC), which replicated a dose previously used to elicit *A. muciniphila* growth in a 10-day feeding intervention (Zhang et al., 2018); (2) an intermediate PAC concentration (6mg/25g, or 6PAC), which is comparable to the PAC level from wine supplementation in our polyphenol composition study; (3) a low PAC concentration (3mg/25g, or 3PAC), to test a reduced PAC level for potential modulation of *A. muciniphila*; and (4) a PAC-free vehicle control (**Figure 3A**).

**Figure 3:**
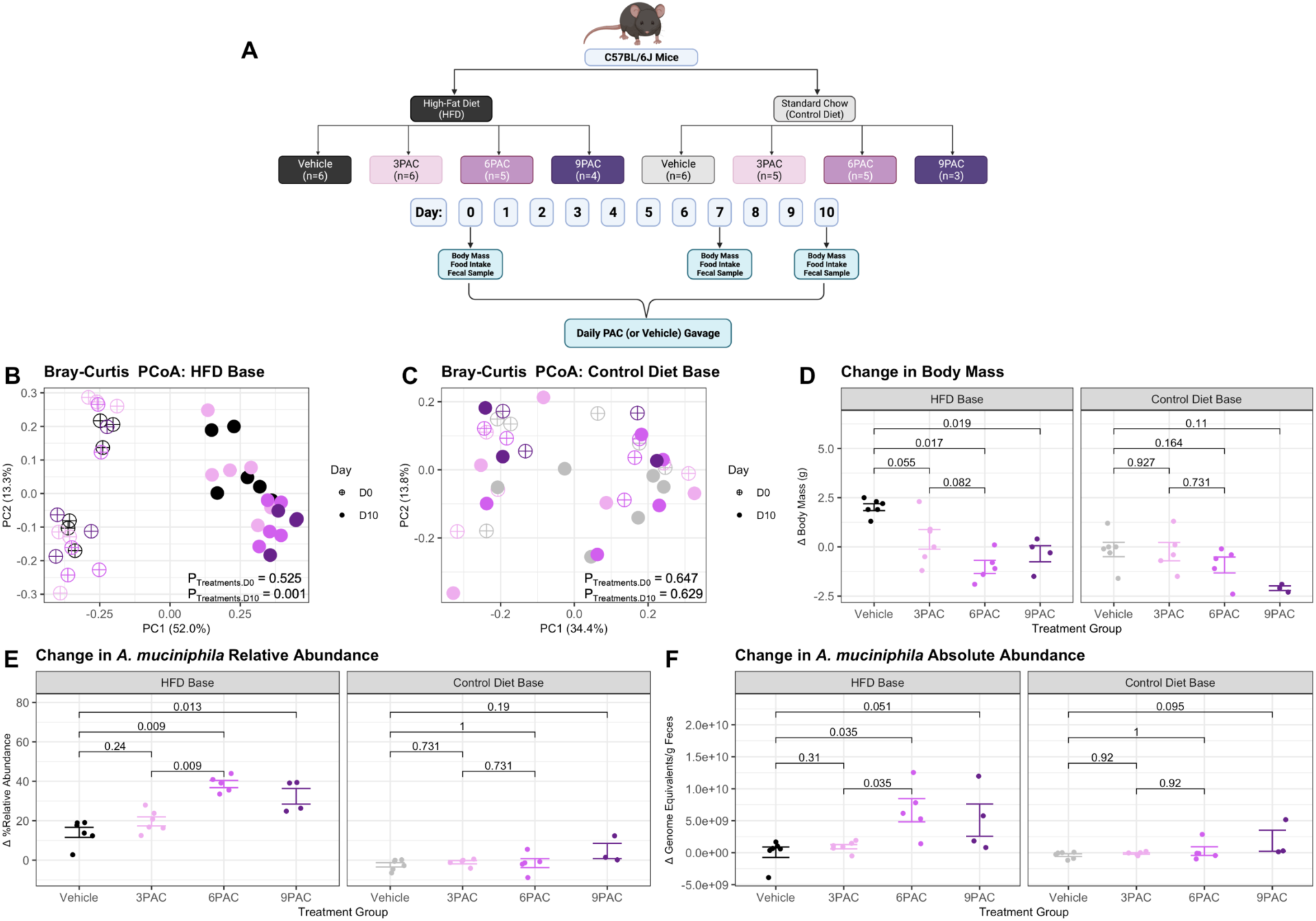
Proanthocyanidin (PAC) supplementation exhibits dose-dependent effects on body mass and microbial community structure under high-fat diet (HFD) but not control diet feeding in mice. **(A)** Adult male C57BL/6J mice consuming either a HFD or control diet base were administered one of the following PAC doses per 25g of body mass over a 10-day intervention: 3mg (3PAC), 6mg (6PAC), 9mg (9PAC), or a PAC-free vehicle control (n=3-6 per condition). PAC doses or vehicle controls were administered daily via oral gavage. **(B)** Bray-Curtis PCoA plot for HFD conditions across Day 0 versus Day 10 of the study. **(C)** Bray-Curtis PCoA plot for control diet conditions across Day 0 versus Day 10 of the study. **(D)** Change in body mass from Day 0 to Day 10 across treatment groups administered each diet base. **(E)** Change in relative abundance of the *A. muciniphila* from Day 0 to Day 10 across treatment groups administered each diet base. **(F)** Change in absolute abundance of the *A. muciniphila* from Day 0 to Day 10 across treatment groups administered each diet base. **(B-C)** Association of samples by treatment group determined at each timepoint using PERMANOVA. **(D-F)** Statistics determined using Mann-Whitney U test with Benjamini-Hochberg correction for false discovery rate.

HFD-fed mice administered either the 6PAC or 9PAC dose exhibited a significant loss of body mass by Day 10, while the 3PAC condition exhibited a marginal but nonsignificant decrease, compared to the vehicle condition (**Figure 3D**). In contrast, mice did not exhibit any significant shifts in body mass under control diet feeding (**Figure 3D**). Gut microbial community composition also differed by treatment group at endpoint in HFD-fed mice (p=0.001, pseudo-F=3.88, PERMANOVA; **Figure 3B**), but not among mice consuming the control diet (p=0.629, pseudo-F=0.868, PERMANOVA; **Figure 3C**). Interestingly, the gut microbial communities of HFD-fed mice administered either 6PAC or 9PAC experienced a relative increase in Bray-Curtis distances relative to baseline, a decrease in alpha-diversity (Shannon index), and lower Firmicutes versus Bacteroidetes relative abundance compared to vehicle controls (**Figure S3A-C**). Moreover, HFD intake induced a decrease in total bacterial abundance in mice administered either the vehicle control or 3PAC supplements, but this bacterial loss was inhibited in the case of 6PAC and 9PAC dosing (**Figure S3D**). These differences in alpha-diversity and beta-diversity, as well as longitudinal maintenance of total absolute abundance, were in part attributable to blooms in *A. muciniphila*. On the HFD, 6PAC and 9PAC dosing induced stark increases in *A. muciniphila* relative abundance (**Figure 3E**). 6PAC dosing also increased *A. muciniphila* absolute abundance, while 9PAC dosing induced a marginal but nonsignificant increase, when compared to the vehicle control (**Figure 3F**). *A. muciniphila* was also detected as an enriched biomarker of 6PAC (p=0.001, coef=1.529, MaAsLin2) and 9PAC (p=0.003, coef=1.501, MaAsLin2) dosing, but the same was not true of 3PAC dosing (p=0.529, coef=0.408, MaAsLin2) or any PAC dosing levels on the control diet by endpoint (p ≥ 0.639, coef ≤ 5.89, MaAsLin2). PAC dosing did not significantly influence gene expression in the proximal colon for our biomarkers of gut inflammation and epithelial integrity under either diet condition (**Figure S3E-L,Q-X**). Overall, we reliably observed changes in gut microbial phenotypes, including expected *A. muciniphila* blooms, among HFD-fed mice supplemented with at least 6mg/25g PAC, while no significant changes were observed at lower PAC doses or under any control diet condition (**Figure 3E-F**; **Figure S3M-P**).

## Discussion

Our data suggest that polyphenols sourced from whole foods can be used to modulate the abundance of *A. muciniphila* in the murine gut microbiota, and show that these effects are amplified against the backdrop of a high-fat, low-fiber diet. By rearing mice for 5 days on raisins, dried cranberries, or dried blueberries, we determined that dried blueberries, which contained the highest TPC and PAC content, were the most efficient in stimulating the growth of *A. muciniphila*. Additionally, through prebiotic supplementation with dealcoholized wine (a polyphenol-rich, PAC-rich food) versus decaffeinated coffee (a polyphenol-rich, PAC-poor food), along with polyphenol-free wine and coffee controls, we established that the effects of polyphenols on *A. muciniphila* growth were most likely attributable to high PAC consumption as opposed to high TPC or other compounds present in the food matrix.

We also determined that a daily mouse dose of 6mg/25g of body mass was sufficient for PAC-induced changes in *A. muciniphila* abundance, which allometrically scales up to a daily dose of 1,265mg PACs for a 65 kg human (Reagan-Shaw et al., 2008). For reference, the Tannat wine used in our polyphenol composition study contains about 200mg of PACs per serving (5oz or 146mL), and one serving of fresh blueberries (1/4 cup or about 40g) contains between 134-249mg of PACs (Gu et al., 2004). Based on these concentrations, consuming approximately two servings of dealcoholized wine and three servings of blueberries would deliver a PAC dose in the range necessary to promote *A. muciniphila* growth, assuming that the necessary dose threshold is similar for humans and mice. Although parallel experiments in humans will be necessary to confirm translatability, and additional work will be required to determine whether common forms of processing, e.g. cooking (Carmody et al., 2019), affects the impact of PAC-rich foods on *A. muciniphila*, our data suggest that *A. muciniphila* abundance might be regulated in humans by selectively ingesting or limiting foods with high PAC content.

Finally, by performing experiments against both a HFD and control diet background, we found that effects of dietary PACs on *A. muciniphila* abundance were amplified on the HFD while the control diet repeatedly mitigated these effects. This finding replicates previous work highlighting the role of isolated PACs in attenuating HFD-induced obesity via blooms in *A. muciniphila* (Anhê et al., 2015; Roopchand et al., 2015) and suggests such effects are achievable via whole food consumption. However, Zhang et al., (2018), which served as the basis for our 9PAC treatment in the PAC titration study, reported increased relative abundance of *A. muciniphila* on a low-fat diet, an effect not observed here whether PACs were diet-derived or delivered in isolated form. It is possible that our inability to replicate these effects was due to the low sample size in our control-9PAC condition, although we also did not detect any effects of control-3PAC or control-6PAC supplementation despite larger sample sizes. It is also possible that differences in baseline gut microbiota profiles across studies led to differential responsiveness of the gut microbiota to PAC supplementation. In the study by Zhang and colleagues, the average baseline relative abundance of *A. muciniphila* among mice consuming a low-fat diet and receiving the 9PAC equivalent dose was 0.79% (Zhang et al., 2018) while mice in our 9PAC treatment group had an average baseline relative abundance of only 0.33%, though the difference between studies did not reach significance (p=0.794, Mann-Whitney U test). Finally, we cannot exclude the possibility that the effects of PAC on *A. muciniphila* abundance depend on a concomitant change in diet that perturbs the gut microbiota and thereby allows for *A. muciniphila* expansion. While this might in principle explain why we observed effects among mice transitioned onto a HFD but not those remaining on a control diet, the findings of Zhang and colleagues suggest that a dietary transition is not a prerequisite for *A. muciniphila* augmentation (Zhang et al., 2018).

That effects of PACs were amplified against the backdrop of a high-fat, low-fiber diet (comprised of 45% of calories from fat, with a fat profile predominantly comprised of long-chain saturated fatty acids) is intriguing because humans that habitually consume a similar diet profile tend to live in industrialized societies (Chadaideh and Carmody, 2021) where both metabolic and neurodegenerative diseases are on the rise (Corbett et al., 2018; Emard et al., 1995) and where *A. muciniphila* is also a prevalent gut commensal (Smits et al., 2017). This suggests that PAC supplementation offers unique leverage in the same environments where *A. muciniphila* modulation could achieve its highest potential health impacts.

While our study did not directly test the consequences of PAC-induced manipulation of *A. muciniphila* for metabolic syndrome or neurodegenerative disease, we determined that interactions between PAC-rich foods and overall diet can reliably influence the abundance of a key member of the gut microbiota that has been repeatedly and, in many cases, causally linked to specific health outcomes. Previous work has shown that blooms of *A. muciniphila* can be maintained with continued PAC treatment, and that early blooms of *A. muciniphila* are predictive of long-term metabolic benefits under a HFD (Anhê et al., 2015; Roopchand et al., 2015). Such data suggest that PAC-induced *A. muciniphila* blooms observed in our study under HFD backgrounds could potentially persist with long-term interventions and confer a protective metabolic effect in humans consuming typical industrialized diets. In contrast, traditional populations such as the Hadza, which are known to consume fiber-rich diets in conjunction with polyphenol-rich foods (Murray et al., 2001), exhibit low levels of *A. muciniphila* in their guts (Smits et al., 2017), which portends that the low level of *A. muciniphila* among our control diet-fed mice may persist regardless of PAC intake. Thus, fiber-rich diets could potentially be a beneficial long-term strategy for mitigating the growth of *A. muciniphila* in Parkinson’s patients, although this would need to be weighed against the parallel possibility that short-chain fatty acids derived from the gut microbial fermentation of dietary fiber promote motor deficits in Parkinson’s disease (Sampson et al., 2016). Taken together, our work supports the promise of PAC-rich whole foods and overall dietary recommendations for regulating *A. muciniphila* abundance depending on individual health needs.

## Acknowledgements

We thank Terence Capellini, Sloan Devlin, Daniel Lieberman, Richard Wrangham, and members of the Carmody lab for constructive feedback. We thank Diana Roopchand for sharing polyphenol-depletion protocols, and Daniel Schrag and Emily Balskus for providing access to essential laboratory equipment. This work was supported by the National Science Foundation (BCS-1919892), the William F. Milton Fund, and the Harvard Dean’s Competitive Fund for Promising Scholarship.

## Author Contributions

K.S.C. and R.N.C. designed the study; K.S.C., K.E.E., and B.E.M. optimized assays; K.S.C. and K.E.E. ran the mouse experiments; K.S.C. and K.E.E. performed 16S rRNA gene sequencing; K.S.C. performed universal 16S qPCR and RT-qPCR; K.S.C. analyzed the data; K.S.C. and R.N.C. wrote the manuscript; all authors contributed to editing the final manuscript.

## Methods

### Animal experiments

All mice were male C57BL/6J purchased from Jackson Laboratory (Bar Harbor, ME, USA) and, unless otherwise noted, cagemates were distributed symmetrically across treatment groups. For the 5-day dried berries experiment, 7-week-old mice were transported in groups of 3 mice per cage, with control chow treatment mice sourced from an in-house colony of age-matched mice. For the 10-day PAC dosage by diet experiment, 8-week-old mice were purchased in groups of 8 mice per cage. For the 12-day polyphenol composition experiment, 7-week-old mice were purchased and transported in groups of 6 mice per cage. Mice were singly housed at the Biological Research Infrastructure (BRI) facility at Harvard University upon arrival and fed a standard low-fat chow (ProLab Isopro RMH 3000: 14.4% of calories from fat; 26.1% from protein; 59.5% from carbohydrate) until their recruitment into the study. All mice were fed *ad libitum*, provided free access to water, and maintained on a 12-hour light-dark cycle. Diets varied based on treatment group and experiment, as described below. Protocols were approved by Harvard University Institutional Care and Use Committee (IACUC 17-06-306).

### Dried berries experiment

For 5 days, C57BL/6J mice were provided *ad libitum* access to either dried blueberries, cranberries, raisins, or a standard chow control diet (n=5-6 per treatment). Fecal samples and body mass measurements were collected daily, during which berry treatments were replenished and food refusals were collected for analysis of food intake. Mice were euthanized on Day 5 of treatment.

### Polyphenol composition experiment

To determine the impact of common whole food polyphenols on *A. muciniphila* abundance, we conducted a 12-day intervention in which mice were supplemented with a daily oral gavage of either wine or coffee as a source of polyphenols. C57BL/6J mice were administered one of the following experimental treatments: alcohol-free wine (wine), alcohol-free and polyphenol-free wine (winePF), caffeine-free coffee (coffee), caffeine-free and polyphenol-free coffee (coffeePF), or a vehicle control, with gavage solutions prepared as described under “Wine and coffee gavage solution preparation” and “Polyvinylpolypyrrolidone (PVPP) removal of polyphenols from food sources” below. Mice were either placed on a high-fat diet (TD.170592: 45% of calories from fat; 15.3% from protein; 39.8% from carbohydrate) or maintained on the standard low-fat chow control diet (ProLab Isopro RMH 3000: 14.4% of calories from fat; 26.1% from protein; 59.5% from carbohydrate) during the 12-day treatment intervention (n=4-6 per condition). Each mouse was measured for adiposity on a baseline and endpoint day. On Day 0, mice in HFD groups were switched onto the HFD chow while the control diet groups continued to receive the standard low-fat chow. For 12 days, beginning on Day 0, mice were gavaged daily with their assigned treatment. Fecal samples were taken on the three days preceding the start of the dietary intervention and on Days 7, 10, and 12 of the dietary intervention. Mice were euthanized on Day 12 of the intervention period.

### Proanthocyanidin (PAC) titration experiment

To understand the effect of proanthocyanidins at more intuitive doses, such as recommended serving sizes of berries or wine, and to evaluate the effect of base diet, mice were assigned to a range of isolated commercial PAC doses. Mice were either placed on the high-fat diet (TD.170592) or maintained on the standard low-fat chow control diet (ProLab Isopro RMH 3000) during the 10-day treatment intervention (n=3-6 per condition). We administered one of the following 4 prebiotic supplements to cohorts of mice consuming either the high-fat or control diet base groups: a maximum dose group (9PAC) receiving the same dose as in Zhang et al. (2018) (9mg PACs/25g mouse), an intermediate dose group (6PAC) receiving 6mg PACs/25g mouse, a dose commensurate with the PACs administered via wine supplementation in our polyphenol composition study, a low dose group (3PAC) receiving 3mg/25g mouse, and a vehicle control group (0PAC) receiving a gavage of 0.5% ethanol in nuclease-free water. Mice received their prescribed PAC dose via daily gavage for the duration of the 10-day intervention based on the protocol below.

### Wine and coffee gavage solution preparation

For the safety of the mice, we used dealcoholized wine and decaffeinated coffee. For the polyphenol-free treatments, polyphenols were removed via the polyvinylpolypyrrolidone (PVPP) removal method discussed under “Polyvinylpolypyrrolidone (PVPP) removal of polyphenols from food sources” below.

Bodega El Porvenir de Cafayate 2018 Amauta Absoluto Tannat wine was used for this study. This vintage was selected because of the high tannin content of its tannat grape varietal. Wine was dealcoholized using a BÜCHI Rotavapor R-300 rotary evaporator (BÜCHI Labortechnik, Flawil, Switzerland). Alcohol removal was confirmed by a 12.5% decrease in mass (the alcohol percentage of the wine). After removing the alcohol, contents of a bottle of wine were homogenized and divided into two halves, with one half destined for polyphenol depletion via the PVPP method discussed below. PVPP treated wine and untreated dealcoholized wine were aliquoted into 50mL tubes and frozen at -80°C at a slight angle. After 24hrs, the tubes were lyophilized for 72hrs to remove the water content from the frozen solutions. Samples were then weighed and reconstituted with deionized water to the desired stock concentration. The concentration of wine was determined based on allometric scaling of a human-equivalent dose of 1 bottle of dealcoholized wine, an upper limit of biological relevance. Dry content in one bottle of wine was calculated based on the density of the dry weight divided by the initial wet weight. The resulting human dose for this wine concentration (0.55g dry wine/kg) was then converted to a mouse-equivalent dose based on the equation for allometric dose translation between humans and mice (Reagan-Shaw et al., 2008): Animal Equivalent Dose (mg/kg) = Animal dose (mg/kg) * (Animal Km ÷ Human Km), where Km is a scaling factor that captures species differences in oxygen utilization, caloric expenditure, basal metabolism, blood volume, circulating plasma proteins, and renal function. The Km values for humans and mice are 37 and 3, respectively (Reagan-Shaw et al., 2008). Applying this allometric translation resulted in a mouse-equivalent dose of 7.34g dry wine/kg mouse, which was reconstituted to a solution of 1.47g/mL. Polyphenol-free wine was less dense (due to the removal of polyphenols), resulting in a stock concentration of 1.30g/mL dry weight to deliver a 0.052g dry weight/kg human-equivalent dose, or 6.52g dry weight/kg mouse-equivalent dose when scaled using the same allometric dose translation method.

Starbucks Decaffeinated Pike Place Roast was used in this study, and was brewed according to package instructions. Briefly, 10g of ground coffee were added to the coffee filter for every 180mL of cold water brewed. The coffee brew was divided into two halves, with one half destined for polyphenol depletion via the PVPP method discussed below. Untreated coffee and PVPP coffee were aliquoted into 50mL tubes and frozen at a slight angle at -80°C. After 24hrs, the tubes were placed into a lyophilizer for 72hrs. Tubes were then weighed and reconstituted with water, reducing the final volume by 22% to maintain comparable phenolic content to between the wine and coffee treatments. To accurately align the total polyphenol dose with the wine condition, deionized water was added to the desired stock concentration of 0.606g dry coffee/mL, which allows for delivery of a 2.97g/kg mouse dose within a 150µL gavage solution for a 30g mouse. The polyphenol-free condition had a stock concentration of 0.497g dry/mL, which allowed for a 2.48g/kg mouse dose. To ensure that mice received an accurate dry-weight dose for their treatment group, gavage solution volumes were calculated daily based on individual body mass. Mice in the vehicle control group received a standard 100µL of solution.

### Polyvinylpolypyrrolidone (PVPP) removal of polyphenols from food sources

In order to control for compounds other than polyphenols in each treatment group, polyphenol-free stocks of wine and coffee were prepared using the polyvinylpolypyrrolidone (PVPP) method of extracting polyphenols (Ranatunge et al., 2017). 5mL syringes were placed into 15mL centrifuge tubes. Cotton wool plugs were placed at the bottom of the 5mL syringes. 0.9g of PVPP powder was then added to the syringe on top of the cotton plug, and powder was compressed using the syringe plunger. 3mL of treatment solution was added to the top of each PVPP column and placed into a centrifuge at 2,000 revolutions per minute (RPM) for 10min. After the centrifuge wash, extracted product in the centrifuge tubes were placed back into the original solution container. 1mL of solution was then added to each PVPP column and centrifuged again at 2,000RPM for 10min. The supernatant was collected into a 50mL tube. This process was repeated 5 more times before disposing of the PVPP columns. This polyphenol depletion protocol successfully removed both TPC and PACs from the wine and coffee substrates (**Figure 2B-C**).

### Measuring total polyphenol content (TPC)

TPC was measured using the Folin-Ciocalteu assay (Ainsworth and Gillespie, 2007). Gallic acid was used to create a dilution series and calibration curve of absorption versus polyphenol content. This curve can be used to plot the absorbance data from the treatment groups to determine accurate measurements of polyphenol content. Treatment group samples were diluted 5-fold by adding Milli Q water to the sample solution. 100µL of sample was placed into 1.5mL tubes, and 100µL of 0.25 N Folin reagent (Sigma-Aldrich) was added to each tube. Each tube was vortexed for 5sec and incubated at room temperature for 3min. 100µL of 1M sodium carbonate was then added to each tube. After tubes were incubated at room temperature for 5min, 700µL of Milli Q water was added, and the tube was vortexed for 5sec. 200µL from each sample tube was placed in a clear, flat-bottomed 96-well plate and the optical density values were read at 726nm on an Epoch 2 BioTek plate reader (BioTek Instruments, Winooski, VT, USA). The standard curve equation was used to determine total polyphenol content of each sample based on absorbance values.

### Measuring proanthocyanidin (PAC) content

PAC content of each sample was measured using the 4-dimethylaminocinnamaldehyde (DMAC) method (Prior et al., 2010). Acidified ethanol was first prepared by adding 12.5mL of hydrochloric acid (Millipore Sigma) to 12.5mL of Milli Q water and 75mL of HPLC grade ethanol (Thermo Fisher). The dilution solution was prepared by adding 80mL of HPLC grade ethanol to 20mL of Milli Q water. DMAC reagent was prepared by adding 0.05g of DMAC (Millipore Sigma) to 50mL of the acidified ethanol solution, which was prepared fresh for each assay. The procyanidin A2 calibration standard was prepared by adding 5mg of procyanidin A2 (Adipogen) to 50mL of HPLC grade ethanol, for a final concentration of 100µg/mL. The procyanidin A2 control sample was prepared by adding 250µL of HPLC-grade ethanol to 1mL of procyanidin A2 calibration standard, for a final quality control sample at 80µg/mL concentration. 32-fold serial dilutions of triplicate stocks were created for the procyanidin A2 calibration standards, the procyanidin A2 control samples, and for each treatment sample. A 32-fold dilution was also prepared for the HPLC ethanol blanks. All samples were diluted using the previously prepared dilution solution. On a clear-bottomed 96-well plate, 210µL of the DMAC reagent solution was added to 70µL of each experimental solution. The 96-well plate was then placed on an Epoch 2 BioTek plate reader (BioTek Instruments, Winooski, VT, USA) and read every minute for 30min at an optical density of 640nm at 25°C. Maximum corrected absorbance readings were calculated by subtracting the average ethanol blank from the maximum absorbance read of each sample. Proanthocyanidin concentrations were determined by calculating the regression equation based on the procyanidin A2 calibration standard dilution series, with triplicate reads averaged for each sample.

### Preparing PAC dosing stocks

Zhang et al. (2018) determined that a 10-day gavage with a dose of 360mg/kg isolated and commercially available proanthocyanidins is sufficient to induce a bloom in *A. muciniphila* (Zhang et al., 2018), which is the equivalent of 9mg of PACs administered daily to a 25g mouse. We decided to match this level and scale down to produce two additional doses for our titration experiment. These additional doses corresponded to 2/3 of the reference dose, or 6mg per 25 g mouse (6PAC), which corresponds to the PAC dose delivered in our wine supplementation experiment, and 1/3 of the reference dose, or 3mg/25g mouse (3PAC). Purified grape seed oligomeric proanthocyanidins (Millipore Sigma) were dissolved in a 0.5% ethanol and Milli-Q water solution. The final gavage stock solution concentrations were the following: 24mg PACs/mL (3PAC), 48mg PACs/mL (6PAC), 72mg PACs/mL (9PAC), and a 0.5% ethanol vehicle control condition (0PAC). Gavage volumes were calculated based on daily body mass measures for each mouse across all PAC treatment groups. Mice administered the vehicle control received a standard 100µL of solution.

### 16S rRNA gene sequencing and analysis

DNA isolation was performed using the PowerSoil DNA Isolation Kit (Qiagen). For each sample, 3µL of purified DNA was then amplified in triplicate via polymerase chain reaction (PCR), using 2µL of forward and reverse barcoded bacterial primers amplifying the V4 region of the 16S rRNA gene (515F and 806R), 1µL of 25mM MgCl2 (New England Biolabs), 10µL of 2.5X 5-Prime Host Mastermix (QuantaBio), and 11µL of nuclease-free water per reaction. Samples were amplified on a Bio-Rad T100 thermocycler (Bio-Rad, Hercules, CA, USA) using the following cycle settings: 94°C for 3min, 35 cycles of 94°C for 45sec, 50°C for 30sec, and 72°C for 90sec, with a final extension at 72°C for 10min. Amplification of pooled PCR products was confirmed using 1.5% gel electrophoresis. Any sample that failed to properly amplify was cleaned using the DNeasy PowerClean Pro Cleanup Kit (Qiagen). PCR products were cleaned on a per-sample basis using AMPure XP beads (Beckman Coulter), then quantified using the Picogreen dsDNA reagent and read with a Spectramax Gemini XS (Molecular Devices Corporation, Sunnyvale, CA, USA) at 480EX/520EM. Amplicon concentrations were determined based on the standard curve, and barcoded amplicons were pooled into a single tube at the calculated volume for 100ng of DNA per sample, with a maximum volume of 30µL per amplicon. The collective pool was further cleaned with Qiaquick MinElute Kit (Qiagen), then loaded onto a mini 1.5% agarose gel and run for 50min at 50V, and sequences of 381bp length were re-extracted using the Qiaquick Gel Extraction kit (Qiagen). The final sample pool was diluted to a 10nM concentration and sequenced on an Illumina HiSeq 2500 (Illumina, San Diego, CA, USA) at the Harvard University Bauer Core Facility. Analysis of 16S rDNA sequencing was performed on the Harvard Odyssey cluster using the QIIME 2 (Quantitative Insights Into Microbial Ecology) software package 2020.2 (Bolyen et al., 2019). Sequences were demultiplexed using the Demux plugin and denoised for quality control using DADA2. Taxonomy was assigned using the Naive Bayes classifier trained on Greengenes 13_8 99% OTUs from the 515F/806R region. Bray-Curtis distances and PERMANOVA comparisons were determined using the beta_group_significance plugin. Differential enrichment analyses were performed using MaAsLin2 (Microbiome Multivariable Association with Linear Models) (Mallick et al., 2021).

### 16S universal qPCR

To estimate gut bacterial density, we performed universal 16S quantitative PCR (qPCR) using the same primer pairs employed for 16S rRNA gene sequencing (515F and 806R). 1µL of template DNA was combined with 12.5µL PerfeCTa SYBR Green SuperMix Reaction Mix (QuantaBio), 6µL nuclease-free H2O, and 2.25µL of each primer. Amplification was performed on a Bio-Rad CFX384 Touch (Bio-Rad, Hercules, CA, USA) in the Bauer Core Facility at Harvard University using the following cycle settings: 95°C for 10min, followed by 40 cycles of 95°C for 15sec, 60°C for 40sec and 72°C for 30sec. Reactions were performed in triplicate with the mean value used in statistical analyses. Cycle-threshold values were standardized against a dilution curve of *Escherichia coli* genomic DNA at the following concentrations (ng/µL): 100, 50, 25, 10, 5, 1, 0.5, plus a no-template (negative) control. Imputed bacterial DNA concentrations were normalized to whole genome equivalents based on a mean genome size of 4.50Mbp, then multiplied by the total template DNA volume (50µL) and divided by the grams of feces utilized in the isolation of template DNA (varied). To calculate absolute abundance of the species *Akkermansia muciniphila* for each sample, total genome copies per gram of fecal matter was multiplied by the relative abundance of *A. muciniphila* determined via 16S rRNA gene sequencing.

### RT-qPCR of proximal colon samples

Using a GentleMACS Octo Dissociator (Miltenyi Biotec, Bergisch Gladbach, Germany), 20-30mg of tissue was dissociated in 700µL of TRK Lysis Buffer (Omega) treated with 2-mercaptoethanol. Tissue lysate was then transferred to a Homogenizer Spin Column (Omega) and spun down for 5min at max speed. Total RNA was extracted from the resulting supernatant using the E.Z.N.A Total RNA Kit I (Omega). DNases were digested from each sample using the DNase I Digestion Set (Omega). Final RNA concentrations were quantified using a Nanodrop One Microvolume UV-Vis spectrophotometer (Thermo Fisher Scientific, Waltham, MA, USA), and RNA extracts were stored at -80°C.

mRNA from each sample was then reverse transcribed to cDNA. 2µg of mRNA were combined with 1µL of random hexamers (Invitrogen) and 1µL of 10mM dNTPs (QuantaBio), then raised to 13µL total volume using nuclease-free water. Samples were heated at 65°C for 5min then chilled on ice for 5min. Next, first-strand synthesis was performed by adding 4µL of First-Strand Buffer (Thermo Fisher), 1µL of DTT (Thermo Fisher), and 1.5µL of SuperScript II (Thermo Fisher). Samples were then incubated using the following settings: 25°C for 7.5min, followed by 42°C for 75min, then 70°C for 15min. Second-strand synthesis was then performed by adding 61.75µL of nuclease-free water, 10µL of NEBuffer2 (New England Biolabs), an additional 3µL of 10mM dNTP (QuantaBio), 5µL of *E. coli* DNA polymerase (New England Biolabs), and 0.25µL of RNaseH (New England Biolabs), then incubating samples at 16°C for 2hrs. Finally, strands were ligated by adding 5µL of *E. coli* DNA ligase buffer (New England Biolabs), 1µL of *E. coli* DNA ligase (New England Biolabs), and another 0.25µL of RNaseH (New England Biolabs), and incubating samples at 16°C for 15min followed by heat-activation at 75°C for 20min. Samples were then cleaned using the PCR Purification Kit (Qiagen) and quantified using a dsDNA HS Assay Kit (Qubit). RT-PCR was then performed using 4µL of dsDNA (diluted to 4ng/µL), which was combined with 10µL of KiCqStart SYBR Green PCR Ready Mix (Millipore Sigma), 1µL of forward primer, 1µL of reverse primer, and 4µL nuclease-free water. The following primer pairs were used to measure gene expression:

IL-6 (forward AAGAAATGATGGATGCTACC, reverse GAGTTTCTGTATCTCTCTGAAG), IL-1β (forward GGATGATGATGATAACCTGC, reverse CATGGAGAATATCACTTGTTGG), Tnfα (forward CTATGTCTCAGCCTCTTCTC, reverse CATTTGGGAACTTCTCATCC), Muc2 (forward AAGAAATGCCCCAATAAC, reverse CATCCATGTAGTGTTCCT), Ocln (forward AAAGCAAGTTAAGGGATCTG, reverse TGGCATCTCTCTAAGGTTTC), Intectin (forward CTCCTCCCTCTTCCCTTT, reverse GCCTTCTGACTTCCACAT), Reg3γ (forward AACAGTGGCCAATATGTATG, reverse TCTCTCTCCACTTCAGAAATC), Pla2g2a (forward GAACAAGAAACCATACCA, reverse ACAACCACTCAAATACAT), Actb (forward GATGTATGAAGGCTTTGGTC, reverse TGTGCACTTTTATTGGTCTC).

Amplification was performed on a Bio-Rad CFX384 Touch (Bio-Rad, Hercules, CA, USA) in the Bauer Core Facility at Harvard University using the following cycle settings: 95°C for 30min, followed by 40 cycles of 95°C for 15sec, 58°C for 30sec and 72°C for 15sec. Reactions were performed in duplicate with the mean value used in statistical analyses. The relative amount of target mRNA was normalized to Actb levels as an endogenous control gene, and data were analyzed according to the 2^-ΔCT^ method.

### Statistical methods

Statistical analyses and data visualization were performed using QIIME 2 (Bolyen et al., 2019) and/or R version 4.1.1 (R Core Team, 2021). Error bars for all plots reflect means ± standard error unless otherwise specified. Student’s t-tests, Mann-Whitney U tests, and MaAsLin2 analyses were corrected using the Benjamini-Hochberg method for false discovery detection.

### Data availability

16S rRNA gene sequencing data have been deposited in the NCBI Sequence Read Archive (accession no. PRJNA763652). Figure source data and additional study data are available upon request.

---

**Figure S1:**
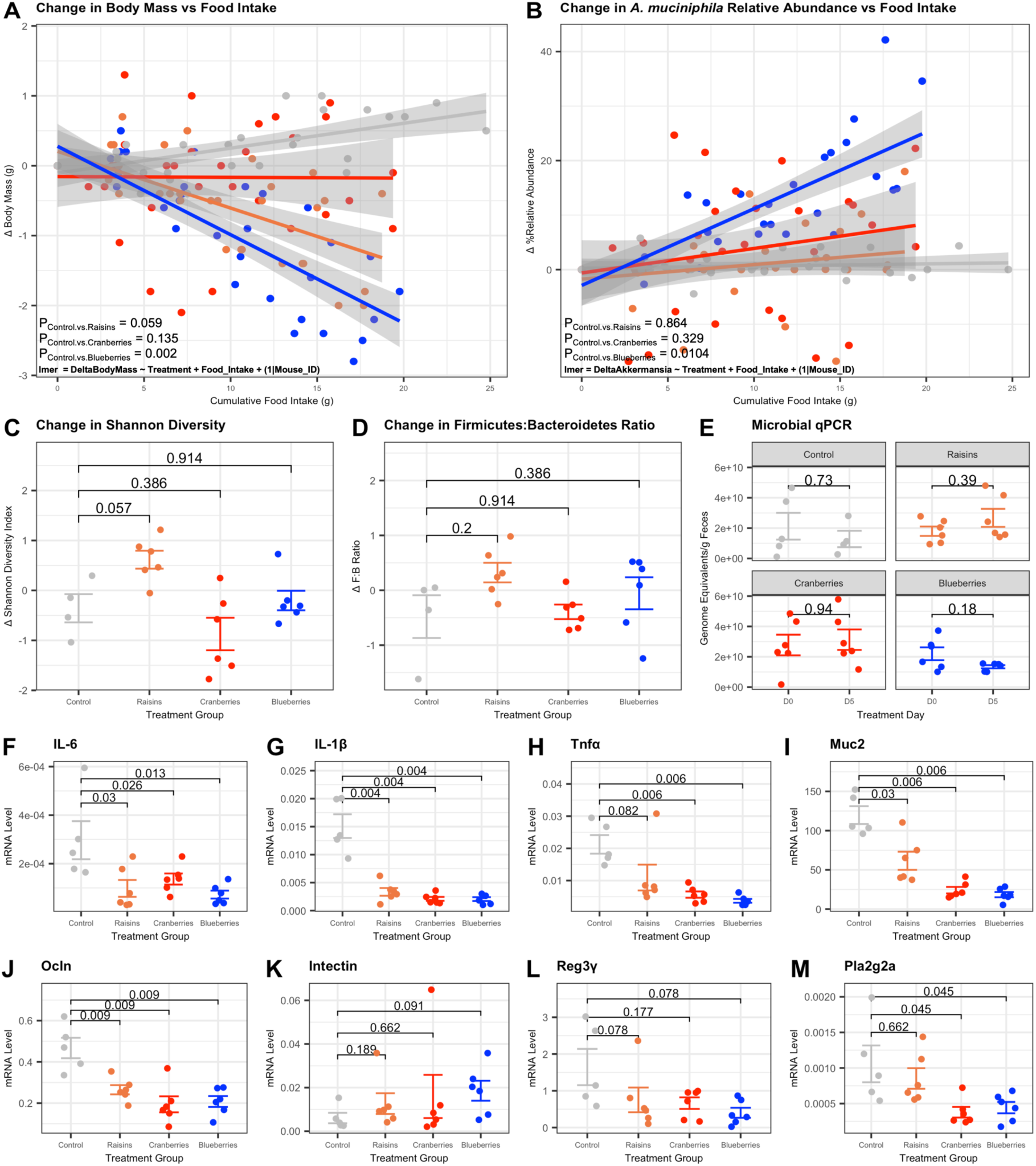
Longitudinal changes in host phenotype and gut microbial community structure across dried berry treatments. **(A)** Change in body mass relative to cumulative food intake across treatment groups. **(B)** Change in the relative abundance of *A. muciniphila* relative to cumulative food intake across treatment groups. **(C)** Change in Shannon Diversity from Day 0 to Day 5. **(D)** Change in the ratio of relative abundance of Firmicutes to Bacteroidetes from Day 0 to Day 5. **(E)** Absolute bacterial abundance on Day 0 versus Day 5, determined using 16S qPCR. **(F-M)** Relative mRNA levels of indicated genes in proximal colon tissue collected from mice receiving each treatment group (tissue control group collected on Day 31 post-intervention, tissue from dried berry treatment groups collected on Day 5 post-intervention). Data represent qPCR of technical duplicates analyzed using the 2^−ΔCT^ method, normalized to expression of the housekeeping gene *Actb*. **(A-B)** Statistics determined using linear mixed effects model. **(C-F)** Statistics determined using Mann-Whitney U test with Benjamini-Hochberg correction for false discovery rate.

**Figure S2:**
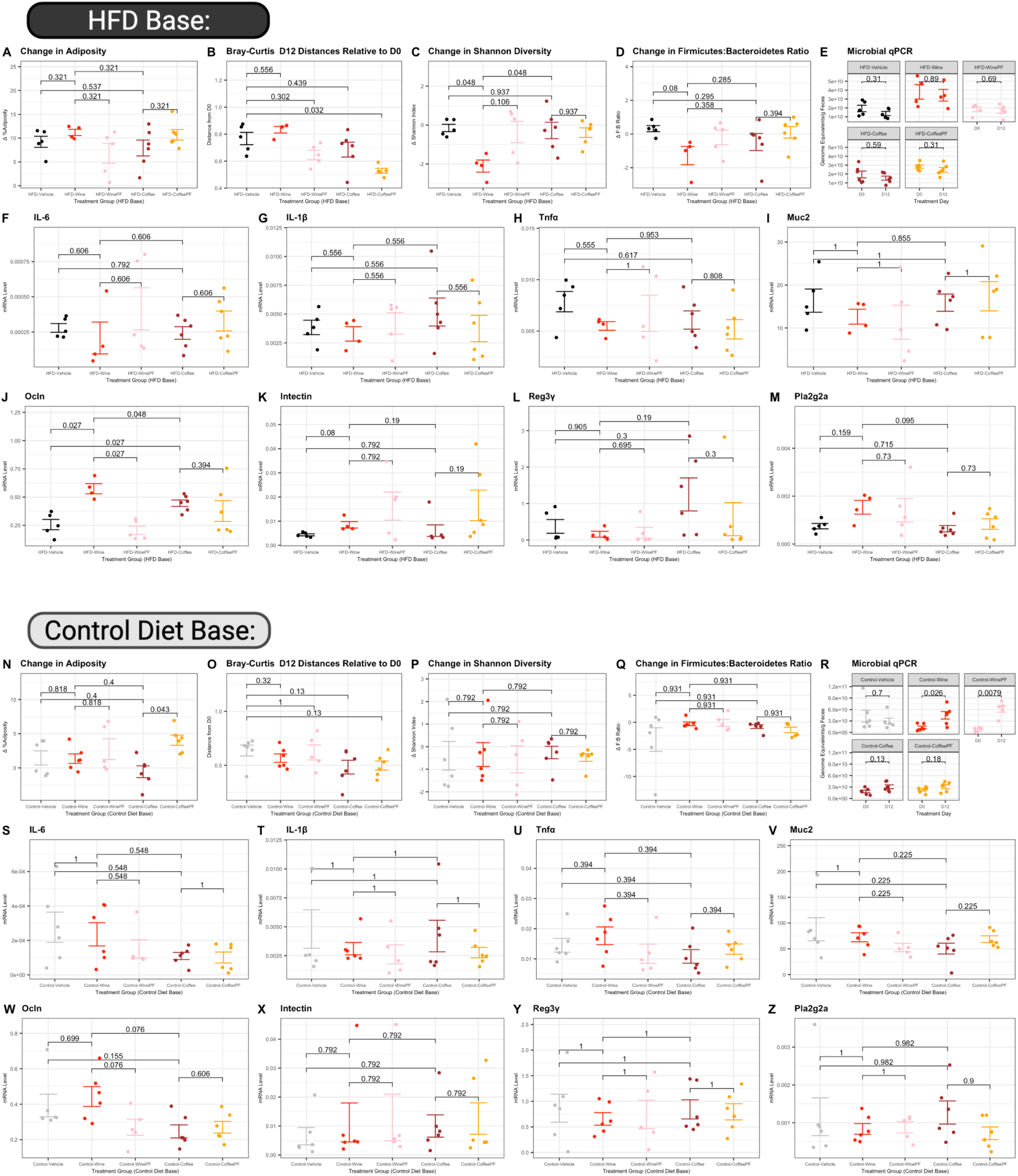
Effects of wine versus coffee supplementation under high-fat diet (HFD) or control diet feeding on host phenotype and microbial community structure. **(A)** Change in percent adiposity from Day 0 to Day 12 across HFD conditions. **(B)** Bray-Curtis distances of fecal samples at Day 12 for mice across HFD conditions relative to their own baseline sample. **(C)** Change in Shannon Diversity from Day 0 to Day 12 across HFD conditions. **(D)** Change in the ratio of relative abundance of Firmicutes to Bacteroidetes from Day 0 to Day 10 across HFD conditions. **(E)** Absolute bacterial abundance on Day 0 vs Day 12 under HFD conditions, determined using 16S qPCR. **(F-M)** Relative mRNA levels of indicated genes in proximal colon tissue collected from mice across HFD conditions (tissue collected on Day 12). Data represent qPCR of technical duplicates analyzed using the 2^−ΔCT^ method, normalized to the housekeeping gene *Actb*. **(N)** Change in percent adiposity from Day 0 to Day 12 across control diet conditions. **(O)** Bray-Curtis distances of fecal samples at Day 12 for mice across control diet conditions relative to their own baseline sample. **(P)** Change in Shannon Diversity from Day 0 to Day 12 across control diet conditions. **(Q)** Change in the ratio of relative abundance of Firmicutes to Bacteroidetes from Day 0 to Day 12 across control diet conditions. **(R)** Absolute bacterial abundance on Day 0 vs Day 12 under control diet conditions, determined using 16S qPCR. **(S-Z)** Relative mRNA levels of indicated genes in proximal colon tissue collected from mice across control diet conditions (tissue collected on Day 12). Data represent qPCR of technical duplicates analyzed using the 2^−ΔCT^ method, normalized to expression of the housekeeping gene *Actb*. **(A-Z)** Statistics determined using Mann-Whitney U test with Benjamini-Hochberg correction for false discovery rate.

**Figure S3:**
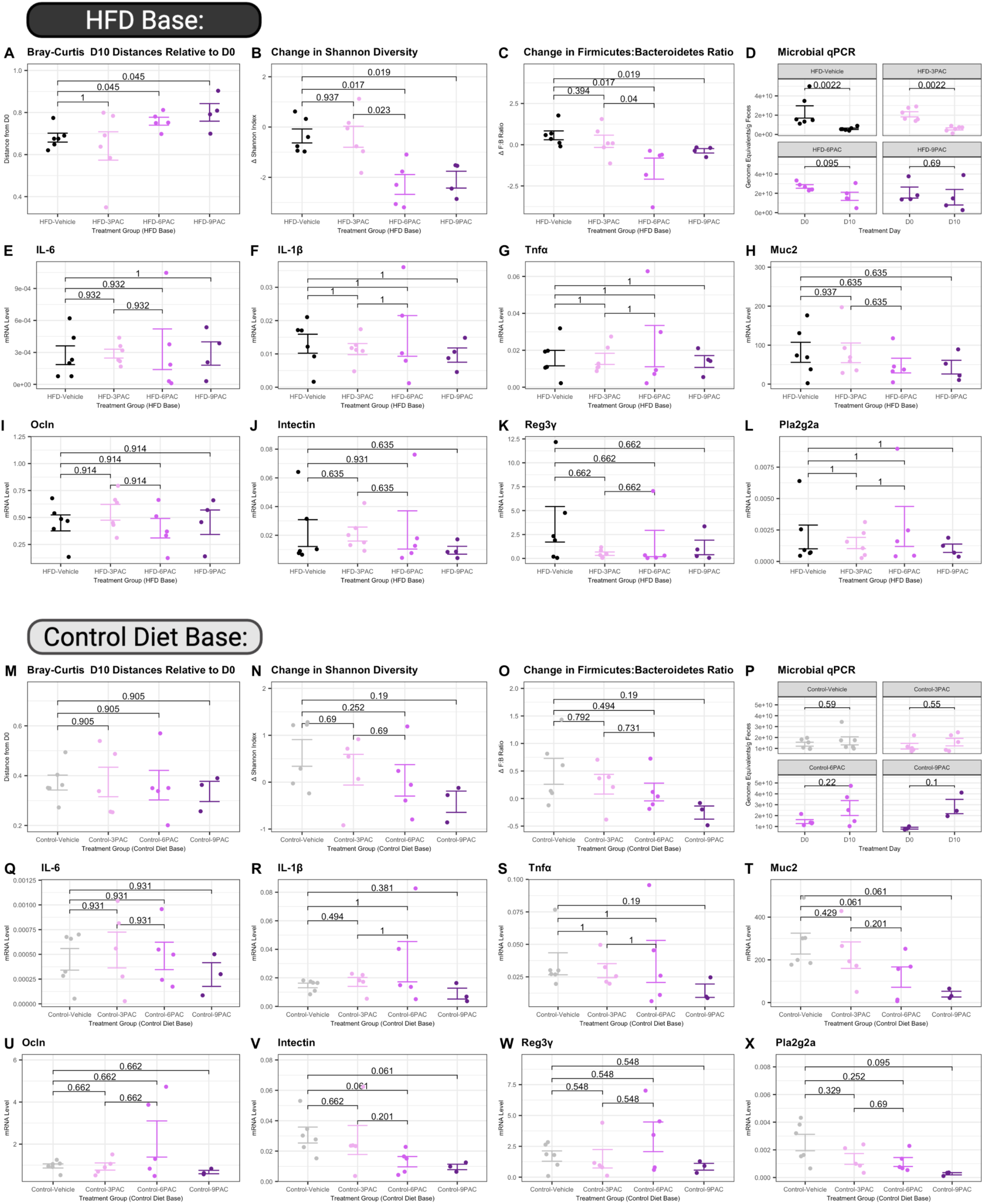
Effects of proanthocyanidin (PAC) supplementation under high-fat diet (HFD) or control diet feeding on host phenotype and gut microbial community structure. **(A)** Bray-Curtis distances of fecal samples at Day 10 for mice across HFD conditions relative to their own baseline sample. **(B)** Change in Shannon Diversity from Day 0 to Day 12 across HFD conditions. **(C)** Change in the ratio of relative abundance of Firmicutes to Bacteroidetes across HFD conditions from Day 0 to Day 10. **(D)** Absolute bacterial abundance on Day 0 vs Day 10 under HFD conditions, determined using 16S qPCR. **(E-L)** Relative mRNA levels of indicated genes in proximal colon tissue collected from mice across HFD conditions (tissue collected on Day 10). Data represent qPCR of technical duplicates analyzed using the 2^−ΔCT^ method, normalized to the housekeeping gene *Actb*. **(M)** Bray-Curtis distances of fecal samples at Day 10 for mice across control diet conditions relative to their own baseline sample. **(N)** Change in Shannon Diversity from Day 0 to Day 10 across control diet conditions. **(O)** Change in the ratio of relative abundance of Firmicutes to Bacteroidetes across control diet conditions from Day 0 to Day 10. **(P)** Absolute bacterial abundance on Day 0 vs Day 10 under control diet conditions, determined using 16S qPCR. **(Q-X)** Relative mRNA levels of indicated genes in proximal colon tissue collected from mice across control diet conditions (tissue collected on Day 10). Data represent qPCR of technical duplicates analyzed using the 2^−ΔCT^ method, normalized to expression of the housekeeping gene *Actb*. **(A-X)** Statistics determined using Mann-Whitney U test with Benjamini-Hochberg correction for false discovery rate.

